# Experimental evidence for antagonistic effects of sperm competition in two sea urchin species

**DOI:** 10.64898/2026.10.08.757654

**Authors:** Luisa Kumpitsch, Lara Maleen Beckmann, Sophie E Hanson, Estefania Paredes, José González Fernández, Lucas Langer, Jan T Lifjeld, Erica H Leder, Jonathan N Havenhand

## Abstract

Sperm competition can occur when sperm from two or more males compete for fertilization of a batch of eggs. While it has been extensively studied in internal fertilizers, it remains poorly understood in external fertilizers. To investigate how sperm competition affects fertilization under sperm limitation, we conducted controlled experiments on two sea urchin species: *Paracentrotus lividus* and *Dendraster excentricus*. Eggs from a single female were exposed to sperm from three individual males separately (non-competitive treatment) or to a pooled sample from three males (competitive treatment). We measured fertilization success as proportion of eggs fertilized after two hours and assessed sperm motility as percentage of motile sperm and curvilinear velocity. For *P. lividus*, we estimated the sperm concentrations where fertilization success was 50% of maximum observed fertilization success (*F_50_*), and the fertilization efficiency of the sperm (*F_e_*), *i.e.* the likelihood that a sperm - egg encounter results in a fertilizing contact, using the fertilization kinetics model of Styan (1998). For *D. excentricus*, fertilization success was measured at a sperm concentration expected to produce 50% fertilization success. Sperm competition reduced fertilization success and increased *F_50_* compared to non-competitive treatments, suggesting antagonistic effects that may compromise fertilization under sperm-limited conditions in external fertilizers.

## 1 Introduction

In external fertilizers, sperm competition - the competition between sperm of two or more males to fertilize a given set of eggs (Wigby & Chapman, 2004) - is a major force shaping reproductive strategies (Parker, 2020; Till-Bottraud et al., 2005). Classical sperm competition theory assumes abundant sperm competing for a fixed number of eggs, following complete fertilization (Bateman, 1948; Parker, 1970), an assumption that often does not hold in external fertilizers, where sperm limitation, polyspermy, and cryptic female choice can lead to incomplete fertilization (Bode & Marshall, 2007; Levitan et al., 2007; Marshall & Evans, 2005; Vogel et al., 1982).

In external fertilizers, the likelihood of fertilization is a function of the ambient sperm concentration (Styan, 1998; Vogel et al., 1982). At low sperm concentrations, eggs may remain unfertilized because sperm are too dilute (Oliver & Babcock, 1992; Pennington, 1985) or fail to penetrate the eggs despite being attracted to them (Marshall & Evans, 2005). Although sperm limitation can favor increased sperm production, selection is expected to favor this strategy only under highly isolated spawning conditions because even low levels of sperm competition can outweigh its benefits (Parker & Lehtonen, 2014). At high sperm concentrations, fertilization success may decline because of polyspermy (Levitan et al., 2007; Styan, 1998) despite the evolution of polyspermy blocks in most taxa, including sea urchins (Longo et al., 1986; Wozniak & Carlson, 2020). Consequently, fertilization is typically maximized at intermediate sperm concentrations where the balance between sperm limitation and polyspermy is optimized (Levitan, 2004; Levitan et al., 2007; Pennington, 1985; Styan, 1998). Within this range, sperm competition could enhance fertilization through cryptic female choice and gamete-recognition mechanisms that favor compatible sperm (Aguirre et al., 2016; Bode & Marshall, 2007; Kosman & Levitan, 2014; Levitan et al., 2007), or reduce fertilization if ejaculates from different males interfere with one another or impair sperm performance (Locatello et al., 2013; Rudolfsen et al., 2015). Sperm motility may further influence these outcomes (Gage et al., 2004; Kupriyanova & Havenhand, 2002). Consequently, the effects of sperm competition on fertilization success likely reflect interactions among sperm concentration, male traits, and female-mediated processes.

Sea urchins are well-established models for studies of external fertilization because spawning can be induced experimentally, allowing controlled fertilization assays (Emlet, 1986; Leuchtenberger et al., 2022; Levitan et al., 2007). The acrosomal protein bindin mediates sperm-egg binding, facilitating species-specific adhesion of sperm to the egg surface (Palumbi, 1999; Vacquier & Moy, 1977; Wessel et al., 2021). At the individual level, variation in the allele coding for bindin can strongly influence intraspecific gamete compatibility (Metz & Palumbi, 1996; Palumbi, 1999). Thus, sperm competition could allow eggs to preferentially interact with genetically compatible sperm, although previous studies found no evidence that prior exposure to heterospecific, low-compatibility sperm reduces fertilization by compatible conspecific sperm (Geyer & Palumbi, 2005; Harper & Hart, 2005; *cf* Lambert, 2000, for tunicates).

We investigated the effects of sperm competition in two species of sea urchins representing two geographically distinct models for examining fertilization dynamics. *Paracentrotus lividus* (Lamarck, 1816) is found in the Mediterranean Sea and European North-Atlantic coast (including Morrocco) and aggregates for spawning (Boudouresque & Verlaque, 2001), creating the possibility of high densities and hence sperm competition. *Dendraster excentricus* (Eschscholtz, 1831), common on the Pacific coast of the USA, can occur in dense aggregations, and many individuals spawn simultaneously causing strong competition among males for fertilizations (Emlet, 1986; see Fig. S1).

Commonly, sperm competition experiments analyze the relative fertilization success of two or more males when their relative sperm contribution can be controlled (see e.g.: Evans & Marshall, 2005; Marshall & Evans, 2005), *i.e.* the sperm is in a “fair raffle” (Parker, 1990). We used two complementary approaches to investigate sperm competition under sperm limitation. In *Paracentrotus lividus*, we investigated fertilization at multiple sperm concentrations, with and without sperm competition, and applied the fertilization kinetics model of Styan (1998) to estimate; i) the sperm concentrations at which fertilization success reaches 50% of maximum observed fertilization success (*F_50_*) and; ii) fertilization efficiency (*Fₑ*), *i.e*. the likelihood that a sperm - egg encounter results in a fertilizing contact. In *Dendraster excentricus*, we compared the proportion of eggs fertilized at a fixed sperm concentration predicted to yield ∼50% fertilization success. By using *F_50_*, or ∼50% fertilization, the “noise” of sperm competition is minimized because the likelihood of polyspermy is reduced. We investigated how sperm competition affects fertilization success and sperm motility by comparing the performance of three males individually (non-competitive treatment) to that of a pooled sample containing sperm from the same three males (competitive treatment) at the same overall sperm concentration. Based on the theoretical framework outlined above, we hypothesized three alternative outcomes: (1) If sperm competition enhances fertilization through increased gamete compatibility or cryptic female choice then fertilization success, fertilization efficiency (*F_e_*), and sperm motility are expected to be higher under competitive conditions, and *F_50_* values should be lower; (2) Alternatively, if sperm competition reduces fertilization through antagonistic interactions among sperm from different males, fertilization success, *F_e_*, and sperm motility are expected to be lower under competitive conditions, and *F_50_* values should be higher; (3) Finally, if treatment has no net effect on fertilization dynamics, no differences between competitive and non-competitive treatments are expected for fertilization success, *F_50_*, *F_e_*, or sperm motility.

## 2 Material and Methods

### 2.1 Sampling and animal maintenance

*Paracentrotus lividus* individuals were collected in October 2022 near the Toralla Marine Station, University of Vigo, Spain (42°12′11.1″N, 8°47′58.2″W). They were maintained for approximately four weeks at 18°C in tanks supplied with filtered (20 µm), flow-through seawater (one tank renewal/day) and fed *Ulva lactuca* regularly. Holding conditions maintained spawning condition beyond the natural spawning peak ending in September (González-Irusta et al., 2010). Experiments were conducted at the Toralla Marine Station in November 2022. *Dendraster excentricus* individuals were collected in April 2023 from East Sound, Orcas Island, Washington, USA (48°41′41.1″N, 122°53′42.8″W). They were maintained for approximately six weeks in tanks supplied with unfiltered, flow-through seawater at Friday Harbor Laboratories, where experiments were conducted in June 2023. Mean seawater temperature during June 2023 was 10.7°C (https://seatemperature.info/june/san-juan-islands-water-temperature.html, 29.10.2025).

### 2.2 Fertilization curves and sperm competition experiments in *P. lividus*

To characterize fertilization success across a range of sperm concentrations commonly used in sea urchin fertilization experiments (Levitan, 2002; Levitan et al., 1991), we performed fertilization assays at six sperm concentrations (see Table 1). Test size (cross-section at largest width and perpendicular length) was measured for each individual, except males from experiment 7, and included as a potential confounding predictor of fertilization success and sperm motility (see Birkhead, 1998; Bode & Marshall, 2007). Spawning was induced by injecting approximately 1 ml of 0.55 M potassium chloride (KCl) into the gonadal region through the mouth and subsequent shaking of the sea urchin (see method in, e.g., Levitan, 2000). Eggs were collected from inverted females into filtered seawater (0.2 µm), whereas “dry” sperm (*i.e.* inactivated sperm before the addition of FSW) was collected from the aboral surface of males and stored on ice for <1 h before use. Individuals were spawned once. Sperm concentration was determined by immobilizing sperm with Lugol’s iodine and averaging the sperm count from twenty small squares in a haemocytometer (Neubauer, Assistent, Germany). Egg concentration was determined by counting eggs within five drops of 50 µl “egg suspension”. Each experiment included one female and three males, each used only once. In each experiment, the eggs of one female were crossed with the three individuals separately (= non-competitive treatment) and mixed (= competitive treatment; see Fig. S2A, B). Procedures were identical save that the competitive treatment required an extra pipetting step since sperm had to be mixed first. Nine experiments were conducted at room temperature (18°C), each with three technical replicates per cross. Fertilizations were performed in 5 ml wells at six sperm concentrations per experiment, with eggs retained in 50 µm mesh filters. After 20 min, excess sperm was removed by rinsing with filtered seawater (0.2 µm). Fertilization success was quantified after 2 h and 24 h by scoring 100 eggs as fertilized or unfertilized. A fertilization curve was fitted for each experiment using the least-squares fitting procedure to the model of Styan (1998).

**Table 1:** Sperm concentrations where fertilization success reaches 50% of maximum observed fertilization success (= *F_50_*) and fertilization efficiency (*F_e_*), i.e. the likelihood that a sperm - egg encounter results in a fertilizing contact (Styan, 1998) in *Paracentrotus lividus*, where the average of three males represents the non-competitive treatment and the mix of those males the competitive treatment. The sperm concentrations (sperm/µl) used to generate the fertilization success curves are shown, where six 10-fold serial dilutions were used from the highest to the lowest concentration. Sperm concentrations for the *F_50_* and *F_e_* were estimated using the fertilization kinetics model in Styan, 1998.

| Experiment | Treatment | Sperm/μl | $F_{50}$ (sperm/μl) | $F_e$ |
| --- | --- | --- | --- | --- |
| 1 | Non-competitive | 0.15 - 15 000 | 871.01 | $1.37 \times 10^{-3}$ |
| 1 | Competitive | 0.15 - 15 000 | 1171.31 | $1.03 \times 10^{-3}$ |
| 2 | Non-competitive | 0.15 - 15 000 | 555.7 | $2.16 \times 10^{-3}$ |
| 2 | Competitive | 0.15 - 15 000 | 2747.87 | $4.22 \times 10^{-4}$ |
| 3 | Non-competitive | 1.5 - 150 000 | 4956.41 | $2.42 \times 10^{-4}$ |
| 3 | Competitive | 1.5 - 150 000 | 11863.24 | $1.02 \times 10^{-4}$ |
| 4 | Non-competitive | 0.75 - 75 000 | 3379.09 | $3.56 \times 10^{-4}$ |
| 4 | Competitive | 0.75 - 75 000 | 13514.13 | $8.64 \times 10^{-5}$ |
| 5 | Non-competitive | 1.5 - 150 000 | 3004.47 | $3.96 \times 10^{-4}$ |
| 5 | Competitive | 1.5 - 150 000 | 12613.99 | $9.58 \times 10^{-5}$ |
| 6 | Non-competitive | 1.5 - 150 000 | 151.65 | $8.79 \times 10^{-3}$ |
| 6 | Competitive | 1.5 - 150 000 | 151.65 | $1.14 \times 10^{-2}$ |
| 7 | Non-competitive | 1.5 - 150 000 | 151.65 | $1.21 \times 10^{-2}$ |
| 7 | Competitive | 1.5 - 150 000 | 151.65 | $6.26 \times 10^{-3}$ |
| 8 | Non-competitive | 1.5 - 150 000 | 151.65 | $8.26 \times 10^{-3}$ |
| 8 | Competitive | 1.5 - 150 000 | 301.8 | $4.41 \times 10^{-3}$ |
| 9 | Non-competitive | 1.5 - 150 000 | 151.65 | $1.05 \times 10^{-2}$ |
| 9 | Competitive | 1.5 - 150 000 | 151.65 | $7.53 \times 10^{-3}$ |

### 2.3 Estimation of fertilization kinetics parameters in *P. lividus*

Across all experiments, fertilization success increased sigmoidally with sperm concentration, with broadly similar response shapes for non-competitive and competitive treatments (see Fig. S3). To quantify treatment effects, we fit fertilization curves for each experimental replicate using the fertilization kinetics model of Styan (1998), which predicts the proportion of monospermic zygotes (*φ_mono_*) as a function of sperm concentration and sperm - egg encounter parameters as described in Equations 1-4:

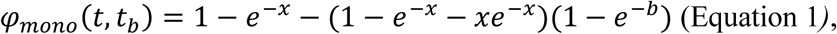

where *x* and *b* can be calculated as:

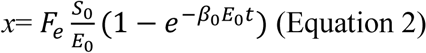

and

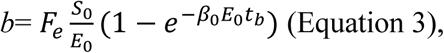

and

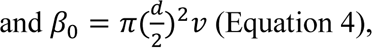

where *φ_mono_*= proportion of monospermic zygotes, *t* = sperm exposure time; *t_b_* = time for the egg to activate a polyspermy block, *x* = the average number of potential fertilizing sperm, *b* = mean number of extra fertilizing sperm that will contact an egg in a time *t_b_* in the population of eggs already contacted once, *F_e_* = fertilization efficiency (the likelihood that a sperm - egg encounter results in a fertilizing contact); *S_0_* = starting ambient sperm concentration (sperm/µl), *E_0_* = initial egg concentration (eggs/µl) and *β_0_*= bimolecular collision rate constant (mm^3^/s). Constants used for all experiments were *E_0_* = 1 egg/µl, *d* = 0.09 mm, *v* = 0.112 mm/s, *t* = 1200 s and *t_b_* = 0.

Constants were chosen according to experimental settings (*E_0_*, *t*), average egg diameter in *P. lividus* (*d*, see Campos et al., 2024), average curvilinear velocity (*v*) and polyspermy (*t_b_*) in the experiments. Fertilization efficiency (*F_e_*) was estimated as the sole free parameter by constrained nonlinear least-squares using the Levenberg-Marquardt algorithm (nlsLM; Levenberg, 1944; Marquardt, 1963), with biologically realistic bounds on *F_e_*. For each fitted curve we computed model predictions across several sperm concentrations (see Table 1 for sperm concentrations, Fig. 1 for model predictions) and obtained *F_50_* numerically as the sperm concentration at which predicted *φ* = 0.5 (*F_50_* was recorded as NA when *φ* did not reach 0.5 within the observed range). We quantified model fit using root mean squared errors (RMSE), a standard measure of predictive error in nonlinear model evaluation (Awasthi et al., 2025; Klimstra & Zehr, 2008; Sedmák & Scheer, 2015), because it expresses the average deviation between observed and predicted fertilization proportions on the response scale (Willmott & Matsuura, 2005). RMSE was calculated for every fertilization curve as in Equation 5:

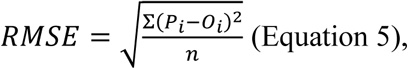

**Figure 1:**
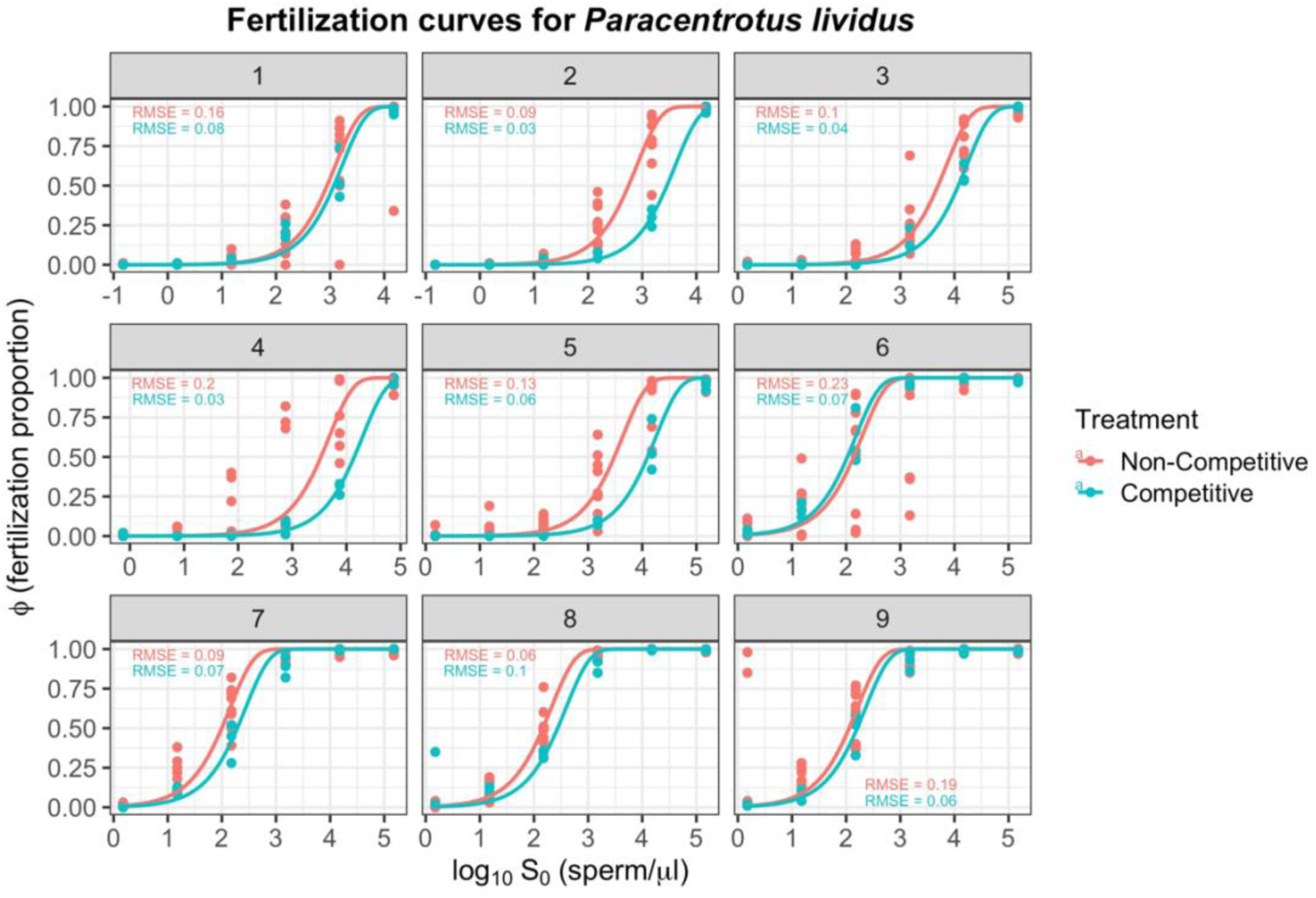
Observed and modelled fertilization curves for each *Paracentrotus lividus* experiment. Panels are faceted by experiment. Points show observed fertilization proportion (*φ*) at each sperm concentration (x-axis plotted as log₁₀ initial ambient sperm concentration, *S₀*). Solid lines are predictions from the Styan (1998) fertilization-kinetics model fitted separately to each replicate. Colors denote treatment: red, non-competitive; blue, competitive. Model constants (egg concentration, egg diameter, sperm speed, exposure time and block time) are as given in Methods. *F_50_* (sperm concentrations where fertilization success reaches 50% of maximum observed fertilization success) and *F_e_* were computed from the model predictions for each fitted curve and used for downstream analyses. Colored RMSE values in each panel show the root mean square error between observed and predicted fertilization proportions for each treatment, where lower values indicate closer agreement between the model and the data. RMSE are on the response scale (= fertilization proportion).

where

*P_i_*= the predicted value for the i^th^ observation;

*O_i_*= the observed value for the i^th^ observation and

*n*= the sample size.

All fitting and data processing were performed with the R-packages *dplyr* (Wickham et al., 2023, v. 1.1.4) and *minpack.lm* (Timur et al., 2022, v. 1.2.4). Predicted curves were plotted with the R-packages *ggplot2* (Wickham, 2016a, v. 3.5.1) and *scales* (Wickham et al., 2011, v. 1.3.0; see Fig. 3).

### 2.4 Fertilization curves in *D. excentricus*

Based on the *P. lividus* results, we first established fertilization curves to identify an intermediate sperm concentration yielding approximately 50% fertilization success for the sperm competition experiments (Fig. S4). This intermediate point maximizes sensitivity in the response curve, and has equal power to detect both increases and decreases in fertilization due to the treatment (Leuchtenberger et al., 2022; Schlegel et al., 2012). Filtered seawater (0.2 µm) was used throughout. Ten males and eight females were randomly spawned and paired for each experiment. Eight experiments were conducted at room temperature (16.3 - 16.4°C). Fertilization assays were performed in 5 ml wells containing eggs at 40 eggs/µl and exposed to sperm concentrations of 1000, 200, 50, 10, or 2 sperm/µl, plus a no-sperm control (Fig. S2C). Procedures followed those described for *P. lividus* (see 2.2), except that eggs were not retained in filters and sperm were immobilized after 25 min by adding 1 ml of 0.55 M KCl (see Leuchtenberger et al., 2022). Fertilization success was assessed after 2 h and 24 h as described in 2.2. An average of 151 ± 45 eggs (mean ± SD) was scored per replicate after 2 h. Mean fertilization success and standard deviations were calculated across all experiments.

### 2.5 Sperm competition experiments in *D. excentricus*

Sperm competition experiments were conducted at the sperm concentration yielding approximately 50% fertilization success (50 sperm/µl; see 2.4, Fig. S4). In each experiment, eggs from one female were fertilized with sperm from three males individually (non-competitive treatment) or with an equal mixture of sperm from the same three males (competitive treatment; Fig. S2D). Fertilizations were performed in 5 ml wells containing 40 eggs/µl and 50 sperm/µl, with an additional no-sperm control. Filtered seawater (0.2 µm) was used throughout. Six experiments were conducted using previously unused individuals. Five experiments included three technical replicates, and one included two. Procedures were identical to those in 2.2. and 2.4. Because fertilization success after 2 h and 24 h was similar (42.4 ± 42% vs. 44.3 ± 40.1%; Fig. S4), only the 2 h data were analyzed. An average of 172.8 ± 44.8 eggs (mean ± SD) was scored per replicate.

### 2.6 Sperm motility

Sperm motility was measured in suspensions prepared from “dry” sperm and filtered seawater. Sperm motility was always measured in individual males (= non-competitive treatment) and for an equal mixture of sperm from the same males (= competitive treatment). For *P. lividus*, “dry” sperm from the males of the experiment was diluted to a final concentration of 10^2^ - 10^3^ sperm/ml. 350 - 450 µL of the sperm dilution was pipetted into the center of an 11/16”-outer diameter O-ring that was placed on a microscope slide. A cover slip was gently placed on top of the O-ring such that it touched the fluid’s surface. The slides and coverslips were coated in bovine serum albumin to reduce the numbers of sperm adhering to the slides or coverslips (Jantzen et al., 2001). Sperm motility was recorded using a high-speed camera (DFK 23UP031 Color Camera, The Imaging Source, 100 Hz) mounted to a microscope (Nikon TE2000S) at 100 x or 200 x magnification. For each male, we filmed ten image sequences (1-2 seconds each). Videos were obtained using NIS software (Nikon, RRID: SCR_014329), and then imported into ImageJ (Version i.52, National Institutes of Health, USA). For all subsequent tracking analyses we used the ImageJ CASA plug-in (Wilson-Leedy & Ingermann, 2007) to track sperm head movement, curvilinear velocity (VCL), and percentage of motile sperm (Alquézar-Baeta et al., 2019). For *D. excentricus*, “dry” sperm was diluted to a final concentration of 10^4^ sperm ml^-1^. 350 - 450 µL of the sperm dilution was pipetted into the center of an 11/16”-outer diameter O-ring, which was glued to a microscope slide using acrylic cement 24 h before filming. We measured sperm motility at 250 x magnification using a high-speed camera (Photron MC1 HSV; at 250 Hz) mounted to a compound microscope (Olympus BH2). For each male, we filmed ten image sequences (1-2 seconds each) and exported these as AVI files with Fastcam Viewer version 3 (Photron Inc., Tokyo, Japan). Subsequent analyses followed the procedures described for *P. lividus* (see Table S1 for used CASA-settings for both species).

### 2.7 Statistical analyses

Analyses were conducted in RStudio (Posit Team, 2024; R Core Team, 2024; R version 4.4.1). For *P. lividus*, mean *F_50_* and *F_e_* were calculated for each non-competitive and competitive treatment within each experiment. To test for differences in *F_50_* and *F_e_* in *P. lividus*, we applied linear mixed-effects models (LME) using restricted maximum likelihood (REML) estimates with the response variables *F_50_* and *F_e_*, a fixed effect treatment (non-competitive *vs* competitive), and two random effects treatments (experiment number and male ID). Male ID accounted for variation in *F_50_* and *F_e_* in an individual male/mix of the males, since we replicated every experiment three times. The normality of model residuals was checked with QQ-plots and homoscedasticity by plotting residuals versus fitted values. The same modelling approach was used for the sperm motility variables, proportion of motile sperm and curvilinear velocity (VCL). Percentage of motile sperm was arcsine square root transformed before statistical analysis to normalize the data.

For *D. excentricus*, means and standard deviation per non-competitive/competitive treatment were calculated for fertilization success, VCL, and percentage of motile sperm for all experiments. Fertilization success and percentage of motile sperm were arcsine square root transformed before statistical analysis to normalize the data. For the remaining analyses, the same procedure described for *P. lividus* was followed.

For *P. lividus*, linear models with the predictor variable test size (*i.e.* body size) and response variables percentage of motile sperm and VCL were used to test for potential effects of test size. Effects of male test size on fertilization parameters (*F_50_* and *F_e_*) in *P. lividus* were tested using generalized linear models with a gamma error distribution and log link since the residuals of the linear model were not normally distributed. The same linear models as described above were used for *D. excentricus* to test for potential effects of test size on fertilization success and sperm motility. To test for effects of VCL and percentage of motile sperm on fertilization success in *D. excentricus*, a linear-mixed effects model using restricted maximum likelihood estimates with the fixed effects percentage of motile sperm and VCL and the random effect experiment number was applied.

To assess whether the benefits of polyandry (= sperm competition; Betters & Levitan, 2018; Evans & Marshall, 2005) depended on variation in male fertilization performance in *D. excentricus*, we calculated the benefit of polyandry for each experiment as the difference in fertilization success between the competitive treatment and either the best-performing male or the mean of the individual males in the non-competitive treatment. Male performance disparity was calculated as the range in fertilization success (maximum - minimum) among individual males within each experiment. We then tested whether male performance disparity predicted the benefit of polyandry using linear regression, with benefit as the response variable and disparity as the predictor. A negative slope would indicate greater benefits when males perform more similarly in fertilization success, whereas a positive slope would indicate greater benefits when males perform more differently.

For data analysis- and organization, we used the R-packages *dplyr* (Wickham et al., 2023, v. 1.1.4), *knitr* (Xie, 2012, v. 1.48), *tidyr* (Wickham et al., 2014, v. 1.3.1), and *tidyverse* (Wickham, 2016b, v. 2.0.0). For the statistical analysis, we applied the packages *lme4* (Bates et al., 2015, v. 1.1.35.5), *car* (Fox & Weisberg, 2019, v. 3.1.3) and *lmerTest* (Kuznetsova et al., 2017, v. 3.1.3) in RStudio (Posit Team, 2024; R Core Team, 2024; R version 4.4.1). All graphs and figures were created using the *cowplot* (Wilke, 2015, v. 1.1.3), *EnvStats* (Millard, 2013, v. 2.8.1), *ggbeeswarm* (Clarke et al., 2016, v. 0.7.2), *ggplot2* (Wickham, 2016a, v. 3.5.1), *ggpubr* (Kassambara, 2016, v. 0.6.0), *hrbrthemes* (Rudis, 2017, v. 0.8.7), *multcompview* (Spencer Graves, Hans-Peter Piepho and Luciano Selzer with help from Sundar Dorai-Raj, 2006, v. 0.1.10), *patchwork* (Pedersen, 2019, v. 1.2.0), *scales* (Wickham et al., 2011, v.1.3.0) and *viridis* (Garnier et al., 2021, v. 0.6.5) packages in RStudio (Posit Team, 2024; R Core Team, 2024; R version 4.4.1), the plugin ScientiFig (Aigouy & Mirouse, 2013) in Fiji (Schindelin et al., 2012) and BioRender (for Fig. S2; created in BioRender. Kumpitsch, L. (2025) https://BioRender.com/ekawckt).

## 3 Results

### 3.1 Fertilization

In both sea urchin species, the competitive treatment negatively affected fertilization outcomes. For *P. lividus*, mean *F_50_* values were lower in the non-competitive treatment than in the competitive treatment (mean ± SD: 1485.92 ± 1813.11 sperm/µl *vs* 4740.81 ± 6013.38 sperm/µl; Fig. 2), corresponding to higher fertilization efficiency (*F_e_*) under non-competitive conditions (0.0049 ± 0.0049 *vs* 0.0035 vs. 0.0042; Table 1, Fig. 2). Treatment (non-competitive *vs* competitive) had a significant effect on *F_50_* (LME, *p*= 0.02, Table 2). Fertilization efficiency (*F_e_*) was strongly negatively associated with *F_50_* (Fig. S5), with higher sperm concentrations required to achieve fertilization in curves with lower fertilization efficiency. For *D. excentricus,* fertilization success was always lower under sperm competition (Figs. 3 and S6, Table S2). Mean fertilization success of individual males in the non-competitive treatment (82.29 ± 21.79%; mean ± SD) was higher than in the competitive treatment from the same males (66.22 ± 27.12%; Fig. 3, Table S2). Treatment (non-competitive *vs.* competitive) had a significant effect on fertilization success (LME, *p*= 0.04, Table 2).

**Figure 2:**
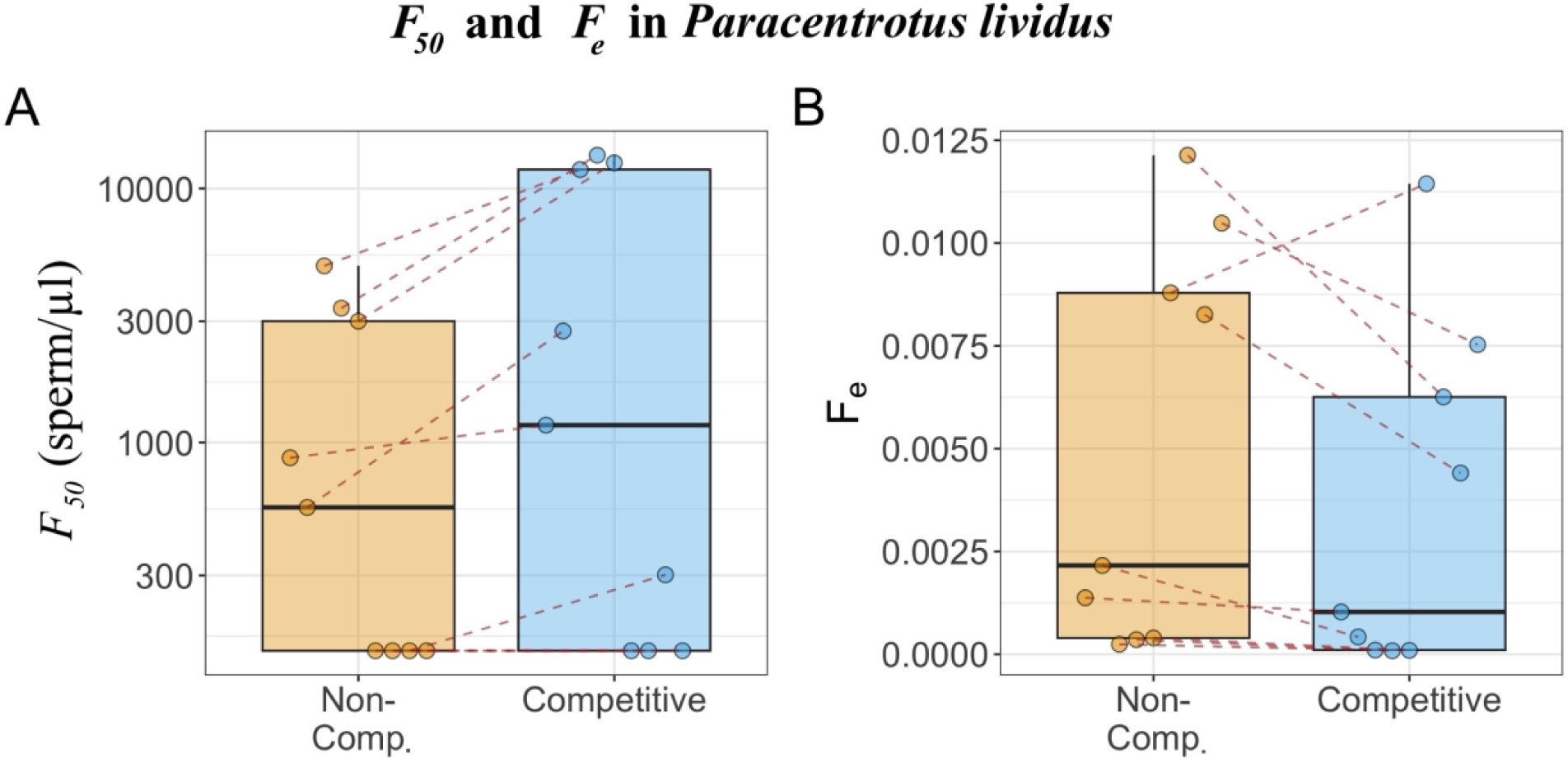
*Paracentrotus lividus*, effects of treatment (non-competitive individual males; “Non-Comp.” and competitive among mixed males; “Competitive”) on (A) *F_50_*, i.e. sperm concentration at which fertilization success reaches 50% of maximum observed fertilization success and (B) fertilization efficiency (*F_e_*) i.e. the likelihood that a sperm - egg encounter results in a fertilizing contact. Boxplots show the median, interquartile range, and range across experiments. Points represent experiment-level estimates, with dashed lines connecting paired non-competitive and competitive treatments within the same experiment. Lower *F_50_* values and higher *F_e_* indicate higher fertilization efficiency.

**Figure 3:**
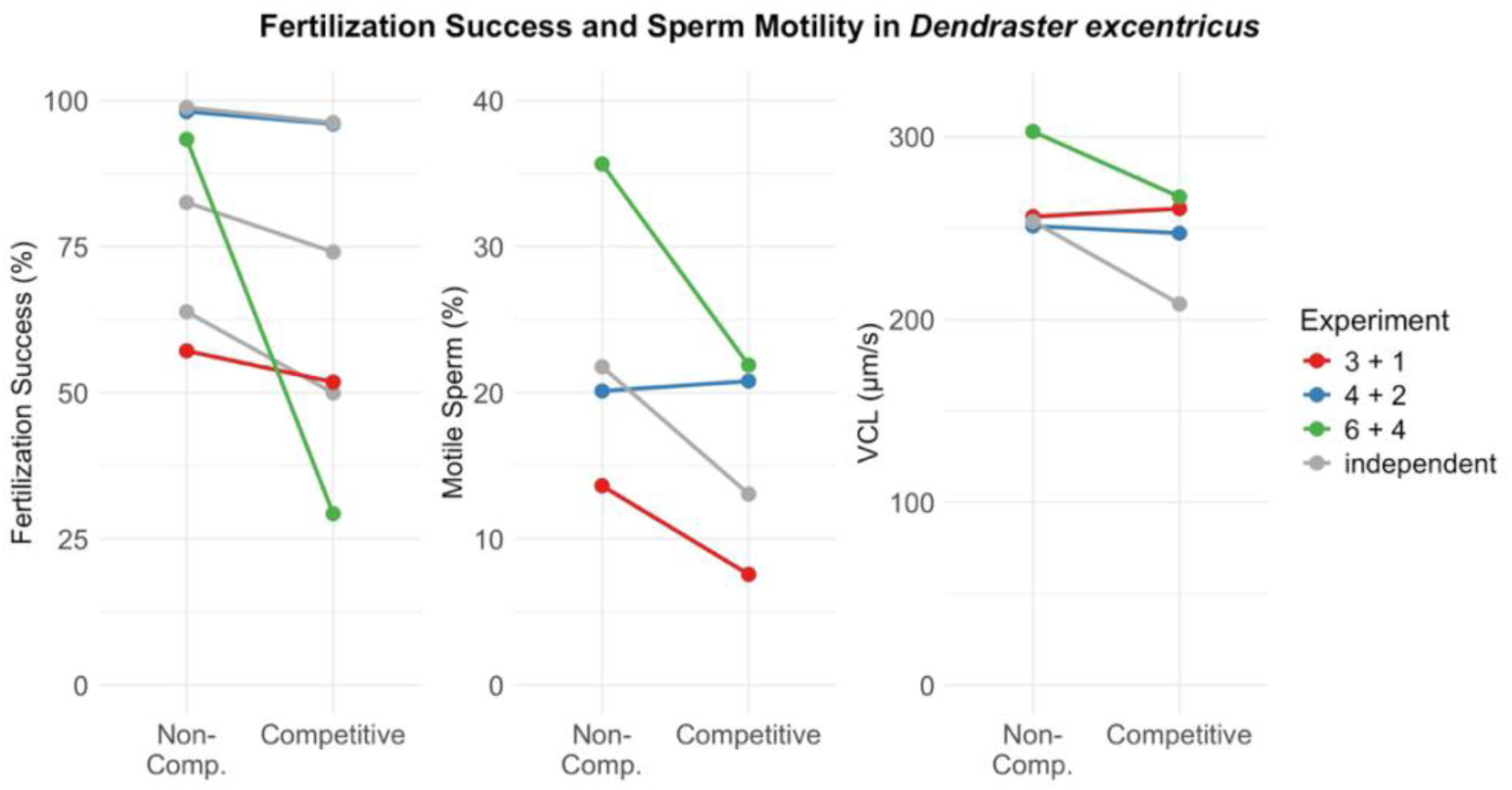
Fertilization success (%), proportion of motile sperm (%), and curvilinear velocity (VCL, µm/s) in *Dendraster excentricus*. In each experiment, fertilization success was compared between three males assayed separately (non-competitive; “Non-Comp.”) and a mixture of the same three males (competitive). Each point represents the mean of the three males, and each line represents one experiment. Grey lines indicate experiments in which fertilization success and sperm motility were measured in different male sets, whereas colored lines indicate paired experiments using the same males. For colored lines, the two numbers in the legend identify the paired fertilization success and sperm motility experiments, respectively (see Tables S1–S2). In sperm motility experiment 3, the non-competitive mean includes only males 7 and 9 because videos for male 8 were unavailable; the competitive treatment and corresponding fertilization success experiment included all three males.

**Table 2:** Results of the linear mixed-effects (LME) models of the effect of treatment (non-competitive *vs* competitive; fixed) and experiment number and male ID (random) in *Dendraster excentricus* and *Paracentrotus lividus*. Model estimate, standard error (SE), degrees of freedom (df), *t*-value and *p*-value are shown. Significant *p*-values (*p* < 0.05) are indicated by asterisks.

| Species | Response Variable | Estimate | SE | df | *t* | *p* |
| --- | --- | --- | --- | --- | --- | --- |
| *P. lividus* | $F_{50}$ (sperm/μl) | 2907.36 | 1152.28 | 25.99 | 2.52 | 0.02* |
| *P. lividus* | $F_e$ | -0.003 | 0.003 | 26.03 | -1.19 | 0.24 |
| *P. lividus* | Percentage of motile sperm | $4 \times 10^{-3}$ | 0.03 | 17.73 | 0.14 | 0.89 |
| *P. lividus* | VCL (μm/s) | $8 \times 10^{-4}$ | 0.05 | 17.47 | 0.016 | 0.98 |
| *D. excentricus* | Fertilization success (%) | -0.24 | 0.11 | 17.41 | -2.22 | 0.04* |
| *D. excentricus* | Percentage of motile sperm | -0.08 | 0.10 | 9.91 | -0.81 | 0.44 |
| *D. excentricus* | VCL (μm/s) | -0.04 | 0.04 | 11.92 | -0.87 | 0.31 |

Across all experiments, the Styan (1998) fertilization kinetics model provided a better fit to the observed fertilization data for the competitive treatments in *P. lividus* (RMSE: 0.03-0.1) than for the non-competitive treatment (RMSE: 0.06 - 0.23). RMSE values were interpreted on the fertilization-proportion scale (0 - 1), where lower values indicate closer agreement between observed and predicted fertilization proportions. In most experiments, fertilization proportions were higher in the non-competitive than in the competitive treatment, with differences between treatments most pronounced at intermediate sperm concentrations and more similar at the lowest and highest concentrations (Fig. 1).

Fertilization success in *D. excentricus* varied greatly among males paired with the same female (Fig. S6, Table S2). The “benefit of polyandry (= sperm competition; Betters & Levitan, 2018; Evans & Marshall, 2005), calculated as fertilization success in the competitive treatment minus that of the best-performing male in the non-competitive treatment, tended to decrease as variation in male fertilization success increased (Fig. S7). However, this negative relationship was not significant in a linear regression including sperm competition benefit as the response variable and male performance disparity, measured as the range in fertilization success among males within each experiment, as the predictor (LM, *p* > 0.1; Table S3). Test size did neither affect *F_50_* or *F_e_* (GLM, *p* > 0.1, Table S4) in *P. lividus* nor fertilization success in *D. excentricus* (LM, *p* > 0.1, Table S4).

### 3.2 Sperm Motility

In both sea urchin species, curvilinear velocity (VCL) was similar between the non-competitive and the competitive treatment (Figs. 3 - 4, Table S5), whereas the percentage of motile sperm was lower in the competitive treatment in two-thirds of the experiments (Figs. 3 - 4, Table S5). In *P. lividus*, mean VCL was slightly lower in the non-competitive treatment (110.77 ± 34.59 µm/s, mean ± SD) than in the competitive treatment from the same males (111.53 ± 26.29 µm/s), as well as the percentage of motile sperm (8.09 ± 3 % *vs* 8.42 ± 4.29 %; Fig. 4), although this difference was not statistically significant (LME, *p* > 0.1, Table 2). In *D. excentricus*, mean VCL was higher in the non-competitive treatment (266.94 ± 53.67 µm/s) than in the competitive treatment from the same males (245.96 ± 26.24 µm/s), as well as the percentage of motile sperm (23.63 ± 13.64 % in non-competitive *vs* 15.83 ± 6.76 % in competitive treatment; Fig. 4) although this difference was not statistically significant (LME, *p* > 0.1, Table 2). Test size did not affect VCL and percentage of motile sperm in *P. lividus* (LM, *p* > 0.1, Table S4) but VCL increased with test size (LM, *p*= 0.05, Table S4) in *D. excentricus*. The percentage of motile sperm and VCL did not affect fertilization success in *D. excentricus* (LME, *p* > 0.1, Table S6).

**Figure 4:**
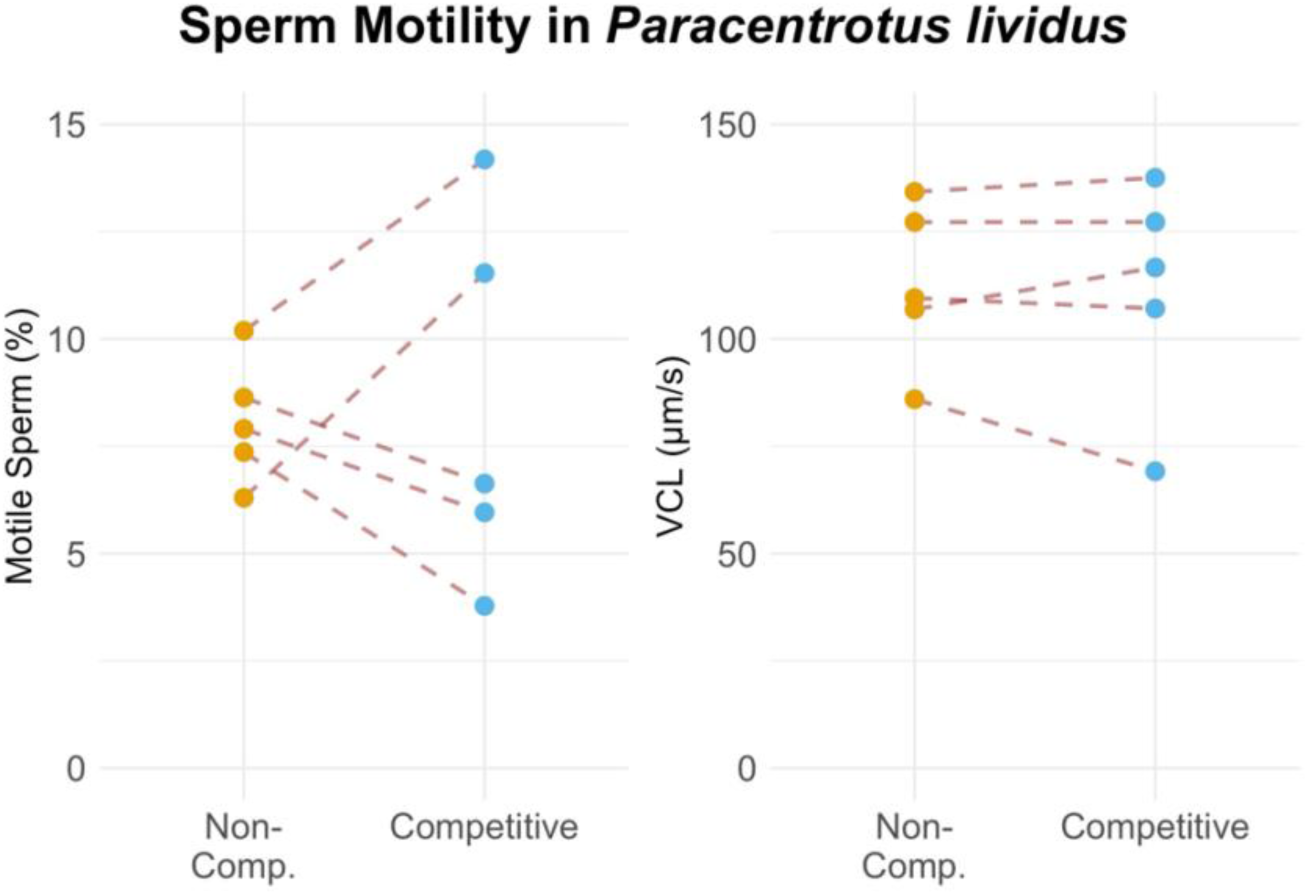
Motile sperm (%) and curvilinear velocity (VCL, µm/s) in *Paracentrotus lividus*, where the average performance of three individual males (= non-competitive treatment; “Non-Comp.”) are compared to that in a mixture of the same three males (= competitive treatment; “Competitive”). Dashed lines connect treatments within each experiment. All experiments were conducted independently, with different males in each experiment.

## 4 Discussion

Sperm competition reduced fertilization performance in both sea urchin species, reflected by lower fertilization efficiency in *Paracentrotus lividus* and lower fertilization success in *Dendraster excentricus*. These findings provide evidence that interactions among rival ejaculates can impose direct fertilization costs, partly supporting our second hypothesis that antagonistic interactions among ejaculates reduce fertilization performance under competitive conditions. Such costs may explain why intense sperm competition is not always favored from the perspective of individual males. In external fertilizers, spawning after a female has released eggs can be costly when rival sperm are already present, and the optimal male - female spawning interval depends on competition intensity, inter-individual distance, and water velocity (Levitan, 2005). Similarly, sperm competition theory predicts reduced sperm expenditure when more than two males spawn synchronously (Parker, 1996). Although both study species form spawning aggregations and likely release gametes synchronously (Boudouresque & Verlaque, 2001; Emlet, 1986; see Fig. S1), our results suggest that males may benefit from reducing direct gamete-level competition when alternative mating opportunities remain available. In the externally fertilizing rainbow darter, for example, males were less likely to spawn when sperm competition intensity was high (Fuller, 1998). However, reduced fertilization performance was not consistent across all *P. lividus* experiments (Fig. 2), potentially reflecting variation in male relatedness or gamete compatibility. Shared alleles at gamete-recognition proteins such as bindin, and male - female genetic compatibility, can maintain high fertilization success in mixed-male sperm environments despite sperm competition (Evans & Marshall, 2005; Levitan & Ferrell, 2006; Palumbi, 1999). A limitation of our study is that sperm in the competitive treatment underwent one additional pipetting step during mixing, which may have imposed minor mechanical stress. Nevertheless, the consistent reduction in fertilization performance across both species suggests that sperm competition can directly reduce fertilization success under the conditions tested.

Comparative studies show that the effects of sperm competition on fertilization success vary markedly among taxa. In the sea urchin *Heliocidaris erythrogramma*, exposing eggs from one female to sperm from two males increased fertilization success relative to a single male (Evans & Marshall, 2005), and in the polychaete *Galeolaria caespitosa* fertilization success overall was higher with sperm from three males than from one (Marshall & Evans, 2005). Those qualitatively different outcomes to our own findings could be partly explained by theory predicting that sperm investment per male is highest when two males compete and declines as the number of competing males increases beyond two (Parker, 1998; Parker et al., 1996). Other studies likewise report benefits of polyandry in sea urchins (*e.g.* Levitan et al., 1992), although these experiments typically included multiple females, increasing the number of male - female combinations and potentially reducing the likelihood of genetic incompatibilities (Evans & Marshall, 2005). In contrast, the effects of sperm competition in *Lytechinus variegatus* depended on male phenotype, resulting in net benefits, costs, or no effect depending on spine colour (Betters & Levitan, 2018). The reduced fertilization success observed in some competitive treatments was attributed to dilution of sperm from the best-performing male. This explanation is unlikely in our study because we would expect the mean of the individual fertilization success to be more similar to that of the mixture, thus there appears to be an additional antagonistic effect within the mixture. Field studies have also reported contrasting outcomes. In *Strongylocentrotus droebachiensis*, synchronous spawning increased fertilization success, although sperm concentration was not controlled for (Pennington, 1985). These studies indicate that the consequences of sperm competition depend strongly on species biology and experimental context. Our results add evidence that antagonistic interactions among ejaculates can reduce fertilization success under controlled sperm concentrations.

Sperm competition may influence fertilization by altering sperm traits and interactions among ejaculates. Males of several externally fertilizing species exhibit adaptive plasticity in sperm traits and spawning behavior in response to sperm competition (Crean & Marshall, 2008; Hadlow et al., 2023; Marshall, 2015). For example, male ascidians (*Styela plicata*) raised at high population density produced sperm with greater size, motility, and longevity than control males (Crean & Marshall, 2008). Consistent with this, sperm mixtures in our study often exhibited a lower proportion of motile sperm than sperm from individual males, suggesting that interactions among ejaculates can reduce sperm performance and thereby contribute to lower fertilization success. Seminal fluid provides one potential mechanism because it modulates sperm motility and chemical signaling involved in sperm-egg interactions (Hopkins et al., 2017). Consistent with this idea, seminal fluid affected sperm velocity and fertilization success in grass gobies, although the direction of these effects depended on male mating tactic: sneaker males benefited from exposure to territorial males’ seminal fluid, whereas territorial males performed worse in sneaker seminal fluid (Locatello et al., 2013). In Arctic charr, fewer sperm cells were initially activated in their own-than in foreign seminal fluid, indicating that sperm activation is under close control of its own seminal fluid (Rudolfsen et al., 2015). In internally fertilizing bees and ants, seminal fluid can protect its own sperm but damage another males’ sperm by incapacitating it (Den Boer et al., 2010). However, whether such interference is efficient or selectively favored remains unclear, because seminal fluid is itself a costly reproductive investment and the fitness benefits of antagonizing rival sperm are uncertain (Hopkins et al., 2019; Ramm, 2020; Simmons & Chan, 2023). These studies suggest that males may influence fertilization outcomes in competitive contexts, with effects that vary depending on species, mating strategy, and social environment.

A key question raised by our results is why sperm competition (polyandry), which can enhance fertilization by increasing opportunities for sperm–egg compatibility (Evans & Marshall, 2005; Marshall & Evans, 2005), instead reduced fertilization success. Possibly, less compatible sperm induce the polyspermy block, preventing more compatible sperm from fertilizing the egg and thereby reducing successful development. In ascidians, even heterospecific sperm can trigger the polyspermy block, inhibiting other (compatible) sperm cells from binding to the egg and hence causing cleavage failure (Lambert, 2000).

Previous studies of the sea urchins *Heliocidaris erythrogramma* and *Lytechinus variegatus* found that the benefits of polyandry increased with differences in individual fertilization success (compatibility) between two males (Betters & Levitan, 2018; Evans & Marshall, 2005). Using a comparable experimental design, we observed the opposite trend. In *D. excentricus*, the effect of sperm competition became more negative as differences in fertilization success among males increased, regardless of whether comparisons were made with the best-performing male (Fig. S7A) or the mean performance of individual males (Fig. S7B), although neither relationship was statistically significant. This pattern is consistent with the hypothesis that less compatible sperm can trigger the polyspermy block and reduce fertilization by more compatible sperm (as in Lambert, 2000). Conversely, when males exhibit similar fertilization success, sperm-egg compatibility may also be more similar, reducing such interference. These findings suggest that the outcome of polyandry depends strongly on sperm-egg compatibility.

## Conclusion

We provide experimental evidence that sperm competition reduces fertilization efficiency in the sea urchins *Dendraster excentricus* and *Paracentrotus lividus*. The observed decrease in fertilization efficiency under competition suggests that males may benefit from behavioral or physiological strategies that minimize direct sperm interference, such as asynchronous spawning or modulation of sperm release. Our results suggest that compatibility between gametes - not just sperm quantity - can shape fertilization success. Future studies should expand this experimental framework to other externally fertilizing taxa to uncover broader patterns in sperm competition, sperm-egg interactions, and fertilization efficiency across marine invertebrates. Ultimately, understanding how sperm competition shapes fertilization success across taxa will provide key insights into the evolution of reproductive strategies in broadcast-spawning marine invertebrates.

## Supporting information

Tables/Figures are marked with S before the numbers

## Data availability statement

Data are available from the SciLifeLab Data Repository: https://doi.org/10.17044/scilifelab.30305140.

## Author contributions

J.N.H. had the original idea for the experiments. L.K., J.N.H., E.H.L., E.P., J.G.F., L.L. and J.T.L. conducted the experiments with *Paracentrotus lividus* at the Toralla Marine Station in Vigo, Spain. L.K., L.M.B., and S.E.H. conducted the experiments with *Dendraster excentricus* at the Friday Harbor Laboratories in Friday Harbor, USA. J.N.H., L.K. and L.M.B. conducted the data analysis, where J.N.H. conducted the modeling. L.K., J.N.H., J.T.L., E.H.L., L.M.B, L.L. and S.E.H. conducted the data interpretation. L.K. led the writing of the original draft of the manuscript. All authors were involved in writing the original draft of the manuscript.

## Funding

This work was supported by the Transnational Access program of the EU H2020-INFRAIA project (No. 871081) AQUACOSM-plus: Network of Leading Ecosystem Scale Experimental AQUAtic MesoCOSM Facilities Connecting Rivers, Lakes, Estuaries and Oceans in Europe and beyond - funded by the European Commission. This study was also supported by the Swedish Research Council VR (2020-05385 to EHL) the FHL Adopt-a-student program, the Rådman och Fru Ernst Colliander Stiftelse, the Herbert och Karin Jacobsons Stiftelse and the Adlerbertska Stiftelse.

## Conflict of interest

The authors have no conflicts of interest to declare.

## Acknowledgements

We thank the staff at the Toralla Marine Station, Vigo, Spain, for assistance with sea urchin sampling, maintenance, and experimental set-up. We also thank Daniel Grünbaum, Richard Emlet, Karen Chan, and Katalin Plummer for valuable discussions, and Friday Harbor Laboratories, Washington, USA, for hosting our experiments.

## Notes

### Competing Interest Statement

The authors have declared no competing interest.

