## Supplementary material for "Experimental evidence for antagonistic effects of sperm competition in two sea urchin species": Tables/Figures are marked with S before the numbers

**Supplementary Data**

Table S1: Parameters used for sperm motility measurements in *Paracentrotus lividus* and *Dendraster excentricus*. Parameters from the table were entered in the ImageJ (Schneider et al., 2012) CASA-plug in (Wilson-Leedy & Ingermann, 2007) to extract curvilinear velocity (VCL; µm/s) and the percentage of motile sperm. For *P. lividus*, microns per 1000 pixels differs depending if sperm motility was recorded at 10x[1] or 20x[2].

| **Sperm motility parameter** | ***Paracentrotus lividus*** | ***Dendraster excentricus*** |
|---|---|---|
| Minimum sperm size (pixels) | 1 | 1 |
| Maximum sperm size (pixels) | 100 | 40 |
| Minimum track length (frames) | 3.3 | 5 |
| Maximum sperm velocity between frames (pixels) | 50 | 8.3 |
| Minimum VSL for motile (µm/s) | 1 | 20 |
| Minimum VAP for motile (µm/s) | 10 | 30 |
| Minimum VCL for motile (µm/s) | 15 | 35 |
| Low VAP speed (µm/s) | 1 | 30 |
| Maximum percentage of path with zero VAP | 1 | 1 |
| Maximum percentage of path with low VAP | 10 | 25 |
| Low VAP speed 2 (µm/s) | 10 | 30 |
| Low VCL speed (µm/s) | 20 | 25 |
| High WOB (percent VAP/VCL) | 40 | 85 |
| High LIN (percent VSL/VAP) | 40 | 65 |
| High WOB two (percent VAP/VCL) | 25 | 85 |
| High LIN two (percent VSL/VAP) | 30 | 65 |
| Frame Rate (frames per second) | 100 | 250 |
| Microns per 1000 pixels | 315[1] (156[2]) | 1087 |

Table S2: Fertilization success (%; mean ± standard deviation SD) in *D. excentricus* across all experiments. The mean ± SD fertilization success is shown for all individual males (= non-competitive treatment) and the mix of the males (= competitive treatment). The replicate number (n) for each treatment is shown.

| Experiment | Male ID | Treatment | Fertilization Success (%) | n |
|---|---|---|---|---|
| 1 | 1 | non-competitive | 77.26 ± 7.34 | 3 |
| 1 | 2 | non-competitive | 48.57 ± 15.46 | 3 |
| 1 | 3 | non-competitive | 65.73 ± 18.39 | 3 |
| 1 | 123 | competitive | 49.96 ± 8.35 | 3 |
| 2 | 4 | non-competitive | 96.97 ± 2.95 | 3 |
| 2 | 5 | non-competitive | 62.99 ± 17.58 | 3 |
| 2 | 6 | non-competitive | 87.64 ± 9.23 | 3 |
| 2 | 456 | competitive | 74.07 ± 12.11 | 3 |
| 3 | 7 | non-competitive | 82.34 ± 16.44 | 3 |
| 3 | 8 | non-competitive | 67.69 ± 37.64 | 3 |
| 3 | 9 | non-competitive | 21.43 ± 4.51 | 3 |
| 3 | 789 | competitive | 51.82 ± 14.25 | 3 |
| 4 | 10 | non-competitive | 100 ± 0 | 3 |
| 4 | 11 | non-competitive | 99.66 ± 0.30 | 3 |
| 4 | 12 | non-competitive | 94.55 ± 4.54 | 3 |
| 4 | 101112 | competitive | 95.96 ± 5.16 | 3 |
| 5 | 16 | non-competitive | 98.73 ± 1.79 | 2 |
| 5 | 17 | non-competitive | 98.96 ± 0.78 | 2 |
| 5 | 18 | non-competitive | 98.65 ± 0.21 | 2 |
| 5 | 161718 | competitive | 96.19 ± 1.85 | 2 |
| 6 | 19 | non-competitive | 97.96 ± 2.04 | 3 |
| 6 | 20 | non-competitive | 98.85 ± 1.99 | 3 |
| 6 | 21 | non-competitive | 83.21 ± 24.62 | 3 |

| | | | | |
|---|---|---|---|---|
| 6 | 192021 | competitive | 29.32 ± 9.68 | 3 |

Table S3: Results of linear regression models testing whether male performance disparity (range in fertilization success among individual males within each experiment) predicts the benefit of polyandry. The benefit of polyandry was calculated either as the fertilization success of the sperm competition treatment (mixed males) minus that of the best-performing individual male (Mix − Best) or as the fertilization success of the sperm competition treatment (mixed males) minus the mean fertilization success of the individual males (Mix − Mean). Reported are parameter estimates estimates, standard errors (SE), t-values, and p-values.

| **Species** | **Response variable** | **Benefit of polyandry** | **Estimate** | **SE** | **t-value** | **p-value** |
|---|---|---|---|---|---|---|
| *D. excentricus* | Fertilization success | Mixed males-Best male | -0.0004 | 0.005 | -0.08 | 0.94 |
| *D. excentricus* | Fertilization success | Mixed males-Mean ind. male | 0.12 | 0.54 | 0.23 | 0.82 |

Table S4: Results of linear models testing the effect of test size on fertilization success, percentage of motile sperm, and curvilinear velocity (VCL) in *Dendraster excentricus* and percentage of motile sperm, and VCL in *Paracentrotus lividus*. Results of generalized linear models show effects of test size on $F_{50}$ and $F_e$ in *P. lividus*. Model estimates, standard errors (SE), *t*-values, and *p*-values are reported for each response variable.

| **Species** | **Response variable** | **Estimate** | **SE** | **t-value** | **p-value** |
|---|---|---|---|---|---|
| *P. lividus* | F50 (%) | 0.02 | 0.09 | 0.20 | 0.84 |
| *P. lividus* | Fe | -0.06 | 0.08 | -0.81 | 0.43 |
| *P. lividus* | Percentage of motile sperm | 0.007 | 0.01 | 0.47 | 0.67 |
| *P. lividus* | VCL (µm/s) | -3.76 | 10.06 | -0.37 | 0.73 |
| *D. excentricus* | Fertilization success (%) | -0.008 | 0.01 | -0.62 | 0.54 |
| *D. excentricus* | Percentage of motile sperm | 0.01 | 0.005 | 2.09 | 0.07 |
| *D. excentricus* | VCL (µm/s) | 3.19 | 1.40 | 2.26 | 0.05 |

Table S5: Curvilinear velocity VCL (μm/s; mean ± SD) and percentage of motile sperm (%; mean ± SD) in *D. excentricus* and *P. lividus* across all experiments. The mean ± SD VCL/percentage of motile sperm is shown for all individual males (= non-competitive treatment) and the mix of the males (= competitive treatment). The replicate number (n) for each treatment is shown.

| Species | Experiment | Male ID | Treatment | VCL (μm/s) | % motile | n |
|---|---|---|---|---|---|---|
| *P. lividus* | 1 | 1 | non-competitive | 106.15 ± 36.15 | 5.45 ± 3.02 | 7 |
| *P. lividus* | 1 | 2 | non-competitive | 121.22 ± 19.09 | 10.55 ± 1.91 | 10 |
| *P. lividus* | 1 | 3 | non-competitive | 108.61 ± 16.15 | 5.56 ± 3.08 | 9 |
| *P. lividus* | 1 | 4 | non-competitive | 91.48 ± 27.83 | 3.63 ± 2.02 | 10 |
| *P. lividus* | 1 | 1,2,3,4 | competitive | 116.64 ± 19.79 | 11.53 ± 5.35 | 9 |
| *P. lividus* | 2 | 5 | non-competitive | 79.98 ± 14.69 | 11.47 ± 5.30 | 7 |
| *P. lividus* | 2 | 6 | non-competitive | 78.16 ± 18.47 | 10.39 ± 1.67 | 9 |
| *P. lividus* | 2 | 7 | non-competitive | 174.96 ± 39.38 | 4.51 ± 1.87 | 10 |
| *P. lividus* | 2 | 8 | non-competitive | 105 ± 37.04 | 5.25 ± 3.56 | 8 |
| *P. lividus* | 2 | 5,6,7,8 | competitive | 107.08 ± 51.62 | 5.96 ± 2.81 | 7 |
| *P. lividus* | 3 | 17 | non-competitive | 82.18 ± 10.07 | 11.67 ± 10.07 | 7 |
| *P. lividus* | 3 | 18 | non-competitive | 99.87 ± 18.14 | 7.74 ± 18.14 | 8 |
| *P. lividus* | 3 | 19 | non-competitive | 199.64 ± 69.34 | 2.67 ± 69.34 | 6 |
| *P. lividus* | 3 | 17,18,19 | competitive | 127.25 ± 44.33 | 3.78 ± 44.33 | 9 |
| *P. lividus* | 4 | X | non-competitive | 145.81 ± 19.45 | 5.95 ± 1.10 | 10 |
| *P. lividus* | 4 | Y | non-competitive | 125.29 ± 4.47 | 9.84 ± 5.90 | 3 |
| *P. lividus* | 4 | Z | non-competitive | 131.71 ± 19.58 | 10.11 ± 4.26 | 8 |
| *P. lividus* | 4 | X,Y,Z | competitive | 137.54 ± 14.32 | 6.63 ± 1.71 | 7 |
| *P. lividus* | 5 | 13 | non-competitive | 93.12 ± 19.10 | 9.67 ± 5.35 | 7 |
| *P. lividus* | 5 | 14 | non-competitive | 92.17 ± 28.18 | 10.06 ± 2.50 | 9 |
| *P. lividus* | 5 | 15 | non-competitive | 91.97 ± 8.69 | 9.09 ± 2.93 | 8 |

| | | | | | | |
|---|---|---|---|---|---|---|
| *P. lividus* | 5 | 16 | non-competitive | 66.61 ± 10.94 | 11.94 ± 0.87 | 5 |
| *P. lividus* | 5 | 13,14,15,16 | competitive | 69.15 ± 8.43 | 14.18 ± 3.53 | 4 |
| *D. excentricus* | 1 | 7 | non-competitive | 267.32 ± 120.36 | 11.81 ± 11.91 | 8 |
| *D. excentricus* | 1 | 9 | non-competitive | 245.37 ± 76.04 | 15.46 ± 8.96 | 10 |
| *D. excentricus* | 1 | 7,8,9 | competitive | 260.78 ± 54.64 | 7.57 ± 6.57 | 10 |
| *D. excentricus* | 2 | 10 | non-competitive | 220.08 ± 80.79 | 8.33 ± 7.61 | 10 |
| *D. excentricus* | 2 | 11 | non-competitive | 268.39 ± 67.54 | 31.96 ± 24.88 | 10 |
| *D. excentricus* | 2 | 12 | non-competitive | 265.25 ± 48.03 | 20.06 ± 20.01 | 10 |
| *D. excentricus* | 2 | 10,11,12 | competitive | 247.29 ± 59.55 | 20.80 ± 14.83 | 10 |
| *D. excentricus* | 3 | 13 | non-competitive | 233.12 ± 28.82 | 23.06 ± 0.67 | 2 |
| *D. excentricus* | 3 | 14 | non-competitive | 184.17 ± 63.15 | 0.89 ± 1.80 | 9 |
| *D. excentricus* | 3 | 15 | non-competitive | 343.83 ± 86.53 | 41.39 ± 25.72 | 10 |
| *D. excentricus* | 3 | 13,14,15 | competitive | 208.60 ± 48.03 | 13.07 ± 11.58 | 9 |
| *D. excentricus* | 4 | 19 | non-competitive | 293.44 ± 172.23 | 29.78 ± 17.81 | 10 |
| *D. excentricus* | 4 | 20 | non-competitive | 371.74 ± 89.32 | 35.50 ± 21.21 | 10 |
| *D. excentricus* | 4 | 21 | non-competitive | 243.63 ± 46.80 | 41.66 ± 14.12 | 10 |
| *D. excentricus* | 4 | 19,20,21 | competitive | 267.16 ± 77.21 | 21.89 ± 19.11 | 10 |

Table S6: Results of a linear-mixed effects model using restricted maximum likelihood estimates with the fixed effects percentage of motile sperm and curvilinear velocity (VCL) and the random effect experiment number. Models were conducted to test for effects of the fixed factors percentage of motile sperm and VCL on fertilization success in *Dendraster excentricus*. Model estimates, standard errors (SE), degrees of freedom (df), *t*-values, and *p*-values are reported for each response variable.

| **Species** | **Fixed Factor** | **Estimate** | **SE** | **df** | **t-value** | **p-value** |
|---|---|---|---|---|---|---|
| *D. excentricus* | Percentage of motile sperm | 1.36 | 0.8 | 2.66 | 1.71 | 0.37 |
| *D. excentricus* | VCL (µm/s) | -0.002 | 0.004 | 2.29 | -0.43 | 0.45 |

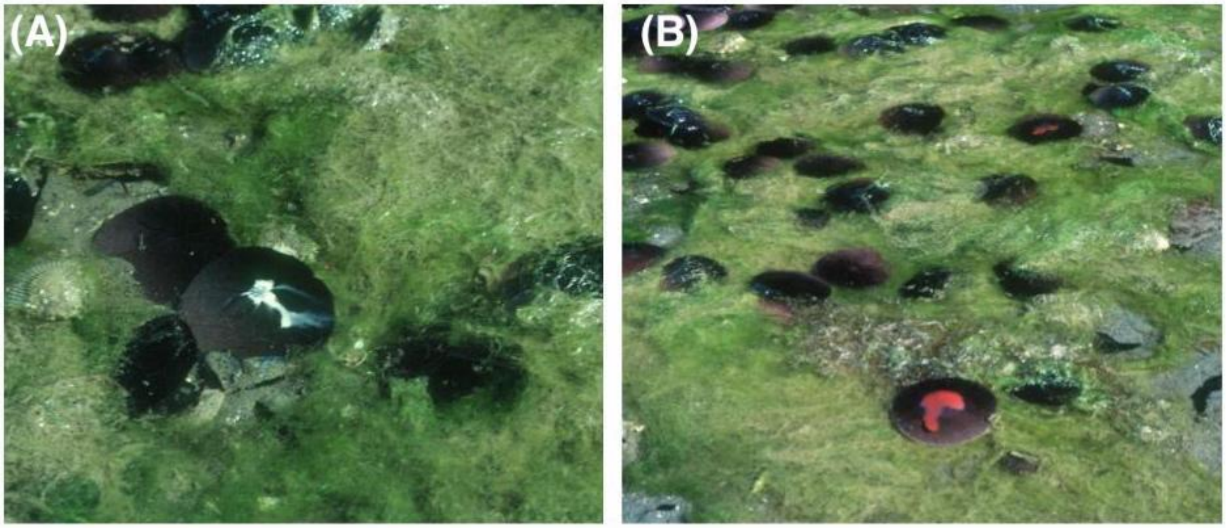


Figure S1: Natural spawning a male (A) and female (B) *Dendraster excentricus* at low tide. Pictures were taken at the south end of Vashon Island (Quartermaster Harbor) in the 1980s. Photographs courtesy of Richard Emlet.

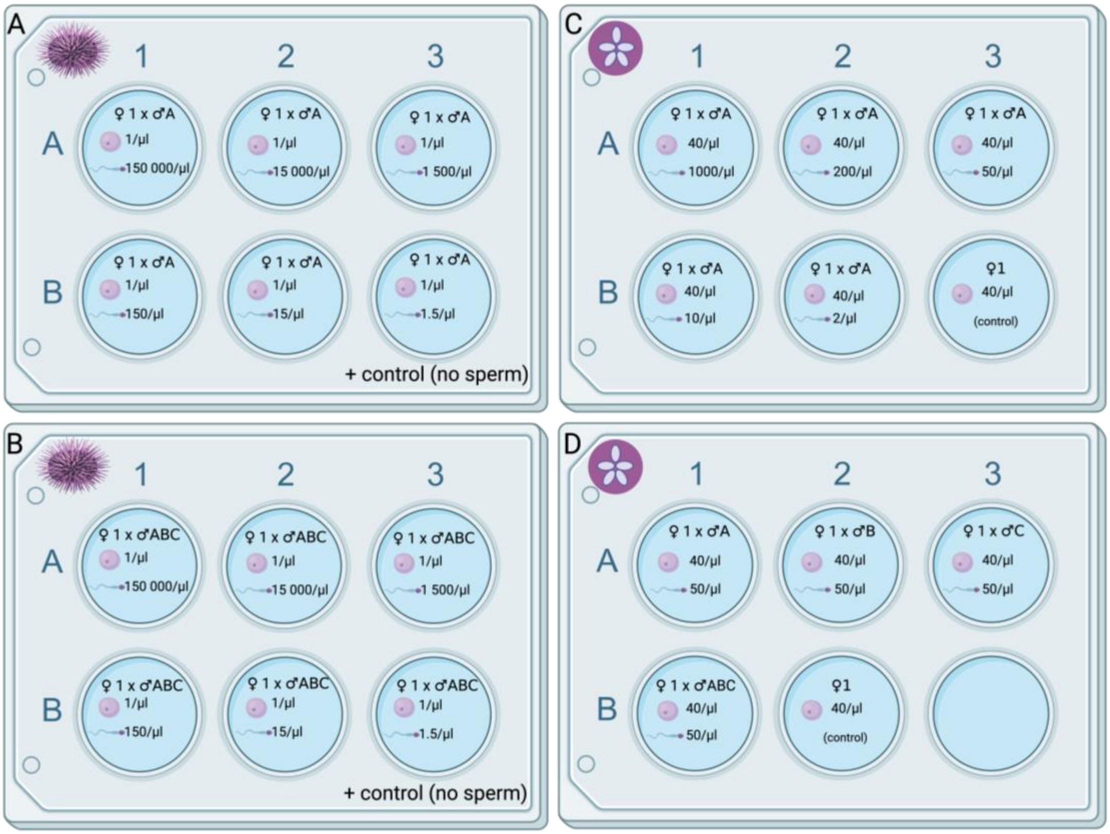


Figure S2: Experimental set-up for fertilization curves (*P. lividus*: A, B; *D. excentricus*: C) and sperm competition experiments (*P. lividus*: A, B; *D. excentricus*: D). The total volume (V) of each well was 5 ml. Egg- and sperm concentrations are shown. Each well plate was replicated three times. For the fertilization curves/sperm competition experiments of *P. lividus* (A, B), egg concentrations were held constant, but sperm concentrations differed among experiments. Sperm concentrations were always 10-fold dilutions (see Table 1), and controls (no sperm) were done in additional wells. Examples of the non-competitive (A) and sperm competition (B) treatment are shown. For the fertilization curves of *D. excentricus* (C), sperm was diluted from 1000 to 2 sperm/µl and egg concentrations were held constant. For the sperm competition experiments, sperm- and egg concentrations were held constant (D).

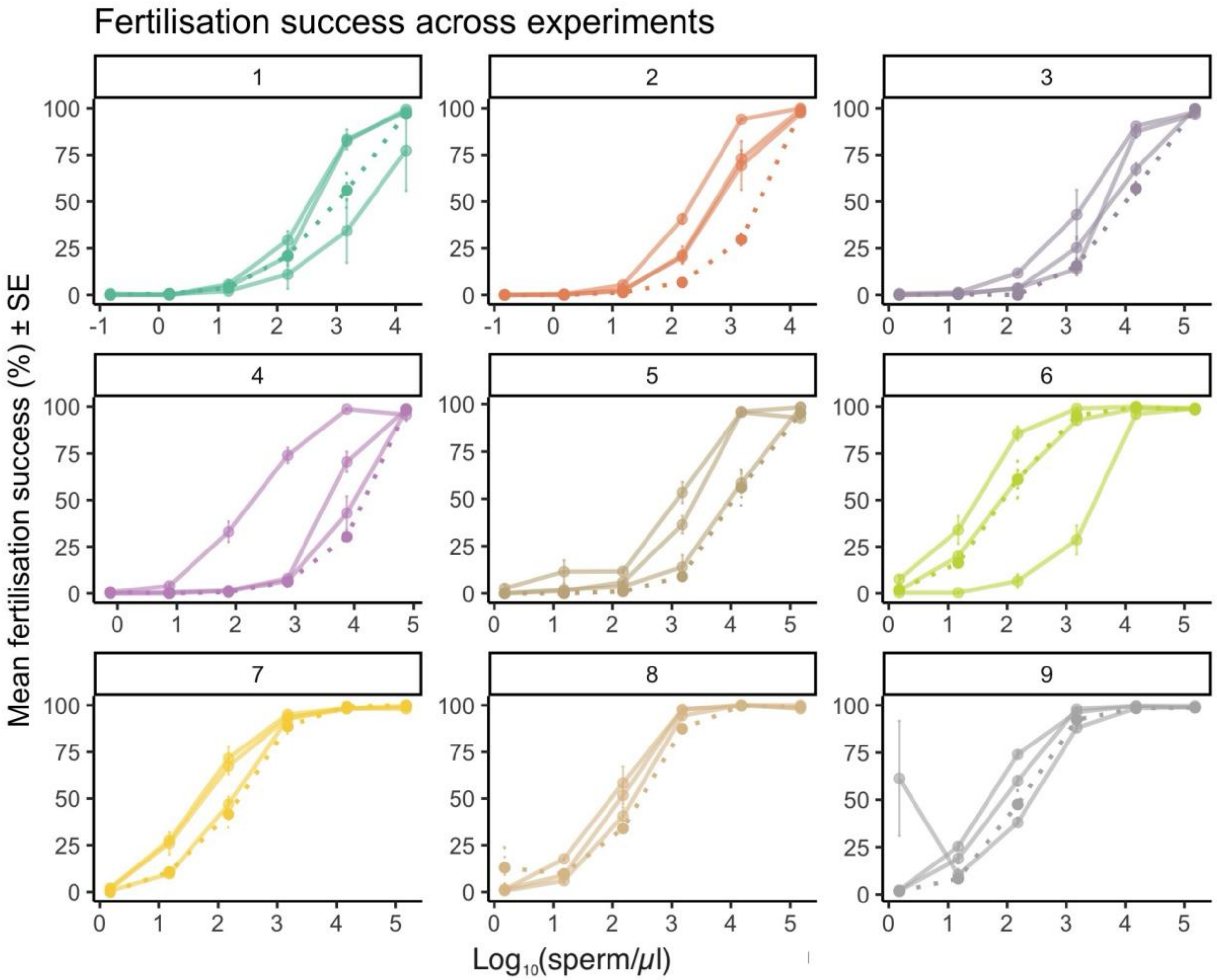


Figure S3: Sperm competition experiments in *P. lividus*, where three individual males were paired with the eggs of a female individually (smooth lines) and as a mixture in the same concentration as the individual males (dashed line). Three replicates were conducted per experiment, whereof the mean is shown here. See experimental procedure in Methods and sperm concentrations in Table 1.

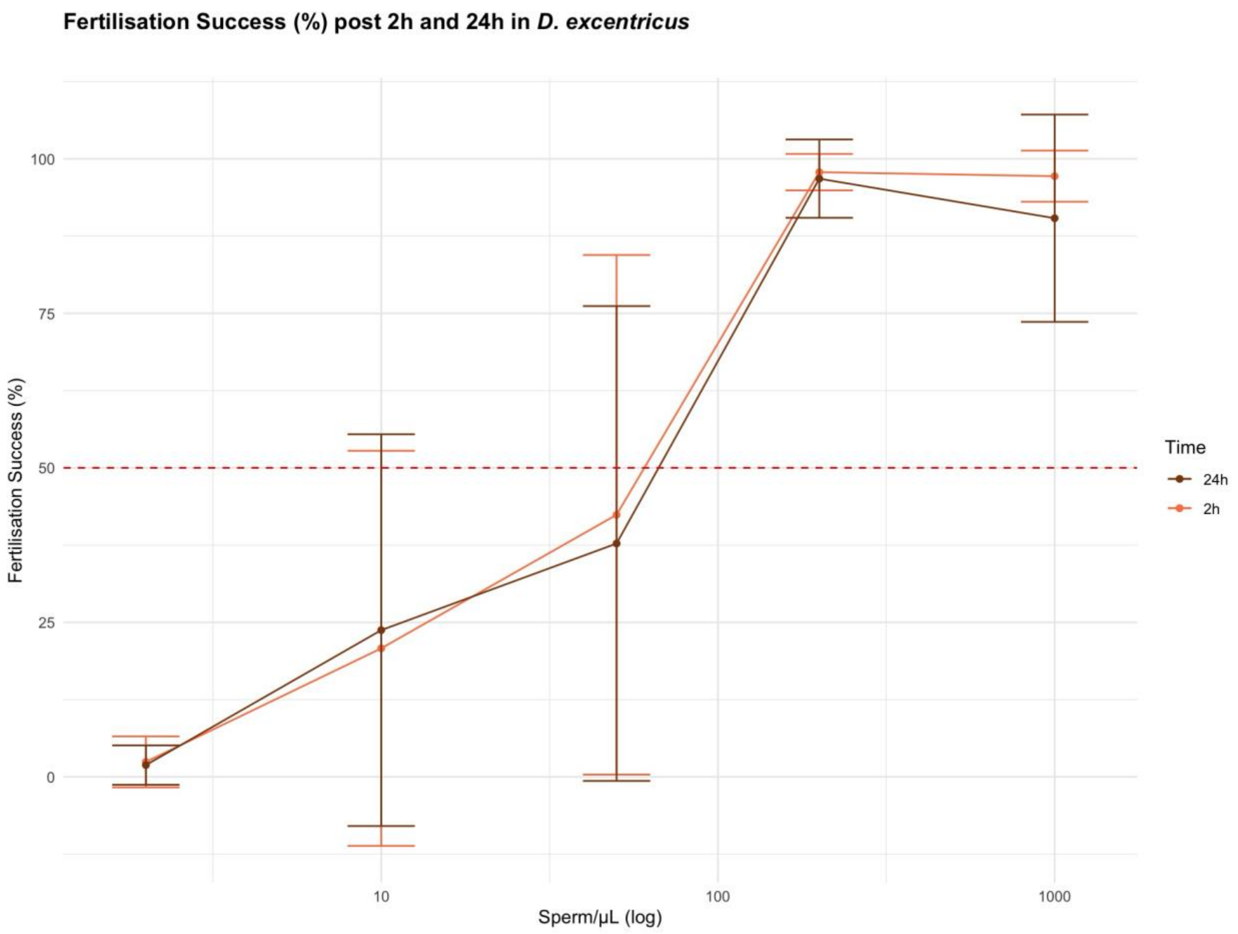


Figure S4: Fertilization success after 2h and 24h *in Dendraster excentricus*. Fertilization success is shown at five different sperm concentrations (log-transformed), where each dot represents the mean, and the vertical bars show the standard deviation at the respective log-transformed sperm concentration. The dotted line marks 50% fertilization success. Note that there are only slight differences between the post 2h- and 24h fertilization success.

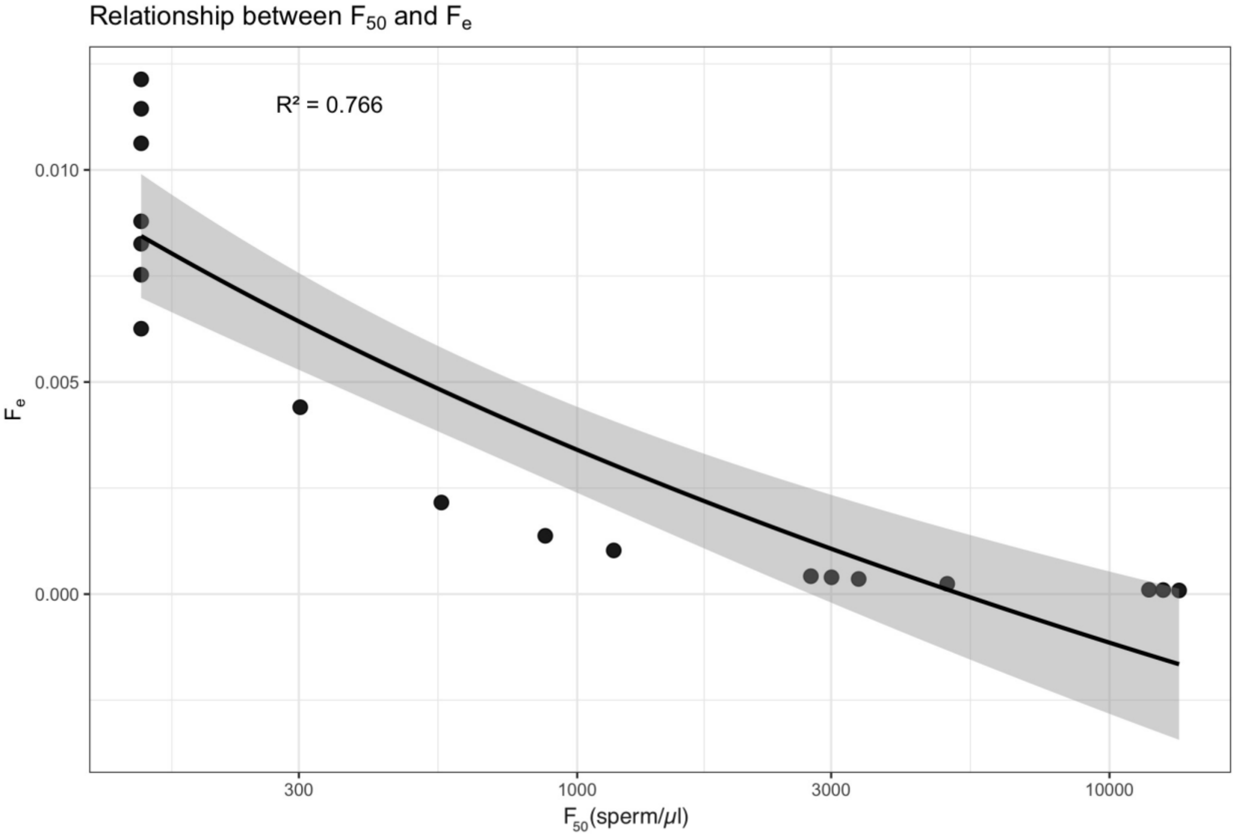


Figure S5: Relationship between fertilization efficiency ($F_e$) and the sperm concentrations where fertilization success reaches 50% of maximum observed fertilization success ($F_{50}$). Each point represents an individual fertilization curve. The solid line shows the fitted linear regression, and the shaded area indicates the 95% confidence interval. The plot shows decrease of fertilization efficiency with increasing sperm concentration.

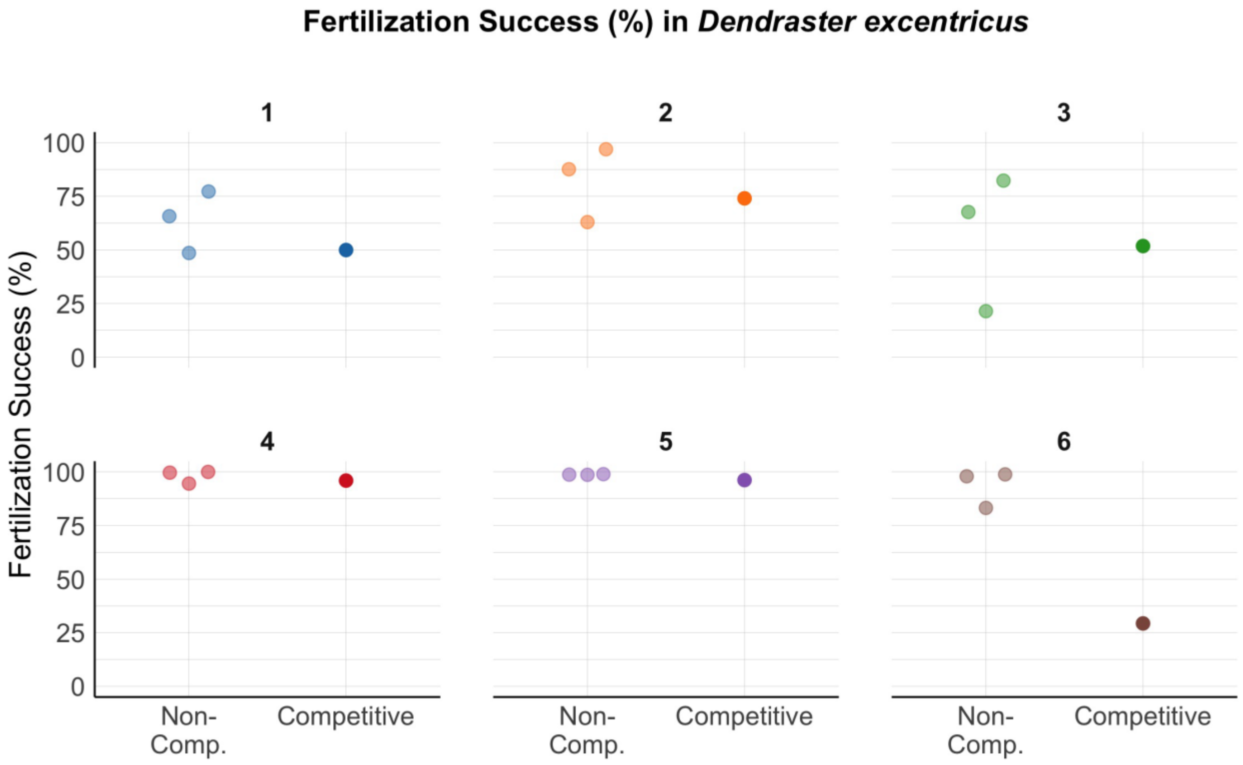


Figure S6: Fertilization success in *Dendraster excentricus*, where individual males (= non-competitive treatment; “Non-Comp.”) are compared to a mix of males (= competitive treatment; “Competitive”). The six experiments conducted are shown. The individuals of the non-competitive treatment are also represented in the competitive treatment, where three individuals make up one mix. A different female was used for every experiment.

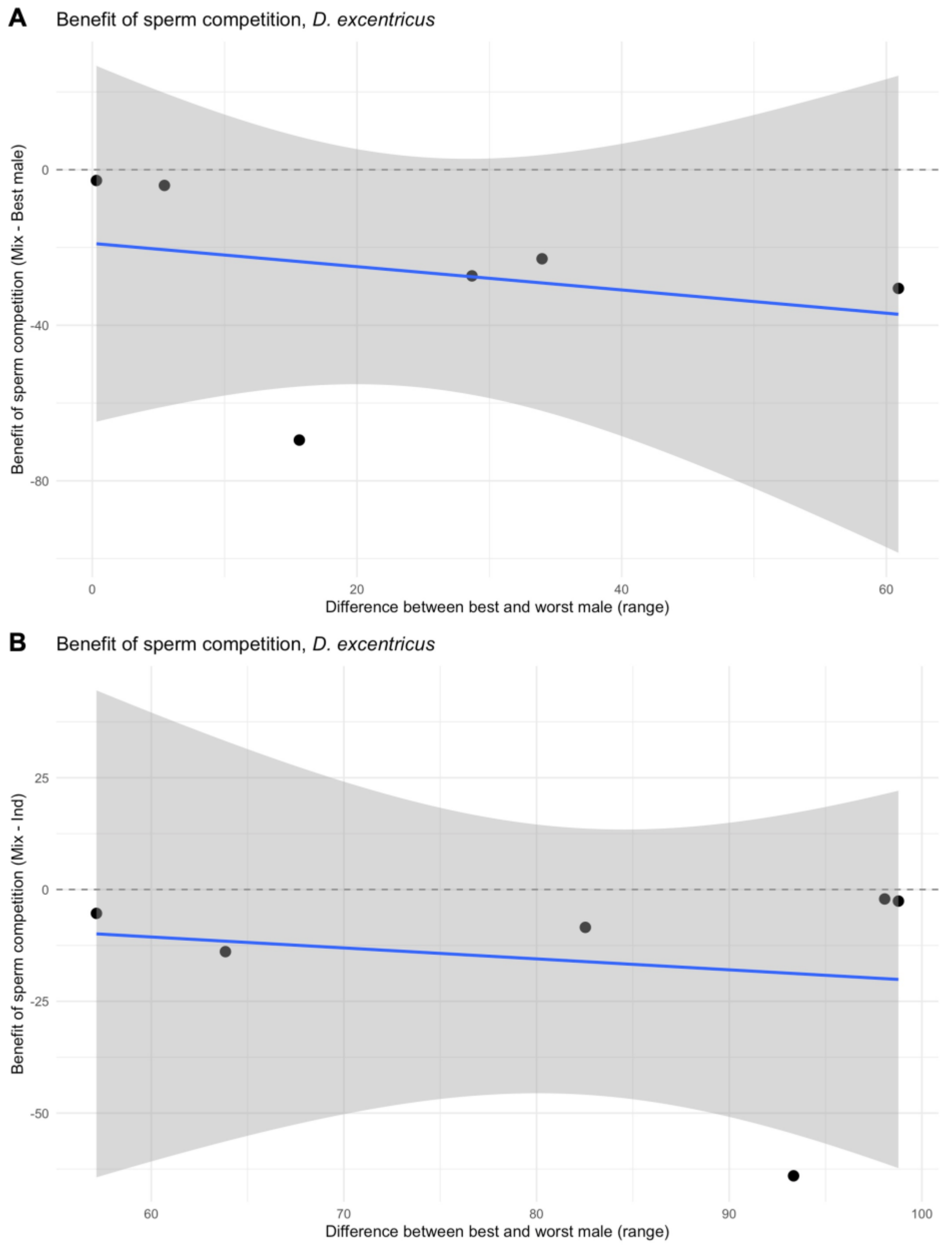


Figure S7: Relationship between male performance disparity and the benefit of sperm competition in *Dendraster excentricus*. The benefit of sperm competition was calculated as fertilization success in the competitive (mixed-male) treatment minus the success of the best individual male within each experiment (A) or minus the average individual success within each experiment (B). Male performance disparity was measured as the range in fertilization success among individual males per experiment. Points represent experiment means, the blue line shows the fitted linear regression, and the shaded area indicates the 95% confidence interval. Values above zero (dashed line) indicate cases where mixed sperm outperformed the best male, while negative values indicate the opposite.
